# Exploring High LET Oxygen-Ion FLASH Radiation for Targeting Pancreatic Cancer in Vitro and In Vivo

**DOI:** 10.1101/2024.12.23.630077

**Authors:** Celine Karle, Domenico Ivan Filosa, Mahdi Akbarpour, Nora Schuhmacher, Stephan Brons, Rainer Cee, Christian Schömers, Stefan Scheloske, Kristoffer Petersson, Thomas Haberer, Amir Abdollahi, Jürgen Debus, Thomas Tessonnie, Mahmoud Moustafa, Andrea Mairani, Ivana Dokic

## Abstract

This study showcases the feasibility of oxygen ion ultra-high dose rate irradiation for in vitro and in vivo experiments at the Heidelberg Ion-Beam Therapy Center. The results indicate comparable effectiveness to standard dose rate in pancreatic cancer cell killing and tumor control, highlighting its potential to address challenges in treating radio-resistant tumors.

## Introduction

Patients diagnosed with pancreatic cancer have the lowest 5-year survival rate of all cancers at around 13%, ranking it as the third most common cause of cancer death [1]. The high mortality rate is attributed to late diagnosis, resulting in advanced tumors growth with unresectable metastases [2]. In the absence of surgery, chemotherapy and external beam radiotherapy represent often the only viable treatment option [3], [4].

Three major challenges reduce the effectiveness of standard radiotherapy in treating pancreatic cancer: proximity to radiosensitive organs at risk (OAR), a hypoxic microenvironment reducing sensitivity to treatment modalities and tumor motion from bowel activity [5], [6].

Addressing these challenges, firstly, pancreatic radiotherapy requires efficient and precisely targeted dose delivery to spare surrounding organs. Particle irradiation offers significantly sharper and more precise dose distributions compared to photon irradiation, allowing superior tumor coverage with minimal exposure to surrounding OAR. In particular, irradiation with carbon ions [7], [8] has demonstrated promising results due to their highly conformal dose distribution [8] combined with high linear energy transfer (LET) [9], [10]. Indeed, studies have shown that high dose-averaged linear energy transfer (LETd) irradiation may be effective in treating pancreatic tumors [11], probably due to the ability to overcome hypoxic tumor resistance by reducing dependence on oxygen levels [9], [12]. High LET irradiation has also been associated with enhanced immune responses [13], providing a dual mechanism for improving treatment outcomes in this context.

To address tumor motion, techniques like breath-hold and abdominal compression have demonstrated promising results [14]. However, ultra-high dose rate (UHDR) irradiation mitigates intra-fractional motion and minimizes motion related uncertainties by delivering doses 100 times faster than conventional radiotherapy. Additionally, the so-called FLASH effect, observed at UHDR, has shown the ability to spare normal tissue while maintaining tumor control, effectively widening the therapeutic window [15], [16]. This approach holds promise for pancreatic cancer treatment, and combined with high LET irradiation, may further enhance the immune response, amplifying its therapeutic potential [17]. However, preclinical studies focusing exclusively on radiotherapy for pancreatic cancer are limited[18], particularly those involving high LET radiotherapy.

Building on the potential of high LET particle therapy at UHDR to address the challenges of pancreatic cancer radiotherapy, this study explored for the first time the effects of oxygen ion UHDR irradiation on three different levels of biological complexity: First, the molecular oxygen consumption in bovine serum albumin (BSA) samples is presented as initial biophysical feasibility study of the setup. Subsequently, in vitro experiments were conducted to evaluate the clonogenic survival of pancreatic cancer cells under hypoxic and normoxic conditions. Thirdly, the UHDR therapeutic efficacy was validated in vivo using the resistant KPC mouse model, a clinically relevant representation of human pancreatic ductal adenocarcinoma (PDAC) [19][20].

## Material and Methods

### Dosimetry

The experiments were conducted at the Heidelberg Ion Beam Therapy Centre (HIT). Details about the facility are given in the supplementary material. For all experiments conducted at UHDR and standard dose rates (SDR) a monoenergetic oxygen ion beam of 325.98 MeV/u was used, incorporating a 3-mm ripple filter at the nozzle to widen the pristine Bragg Peak (BP). Given this default, the experiments were conducted with the setups shown in Figure 1.

**Figure 1:**
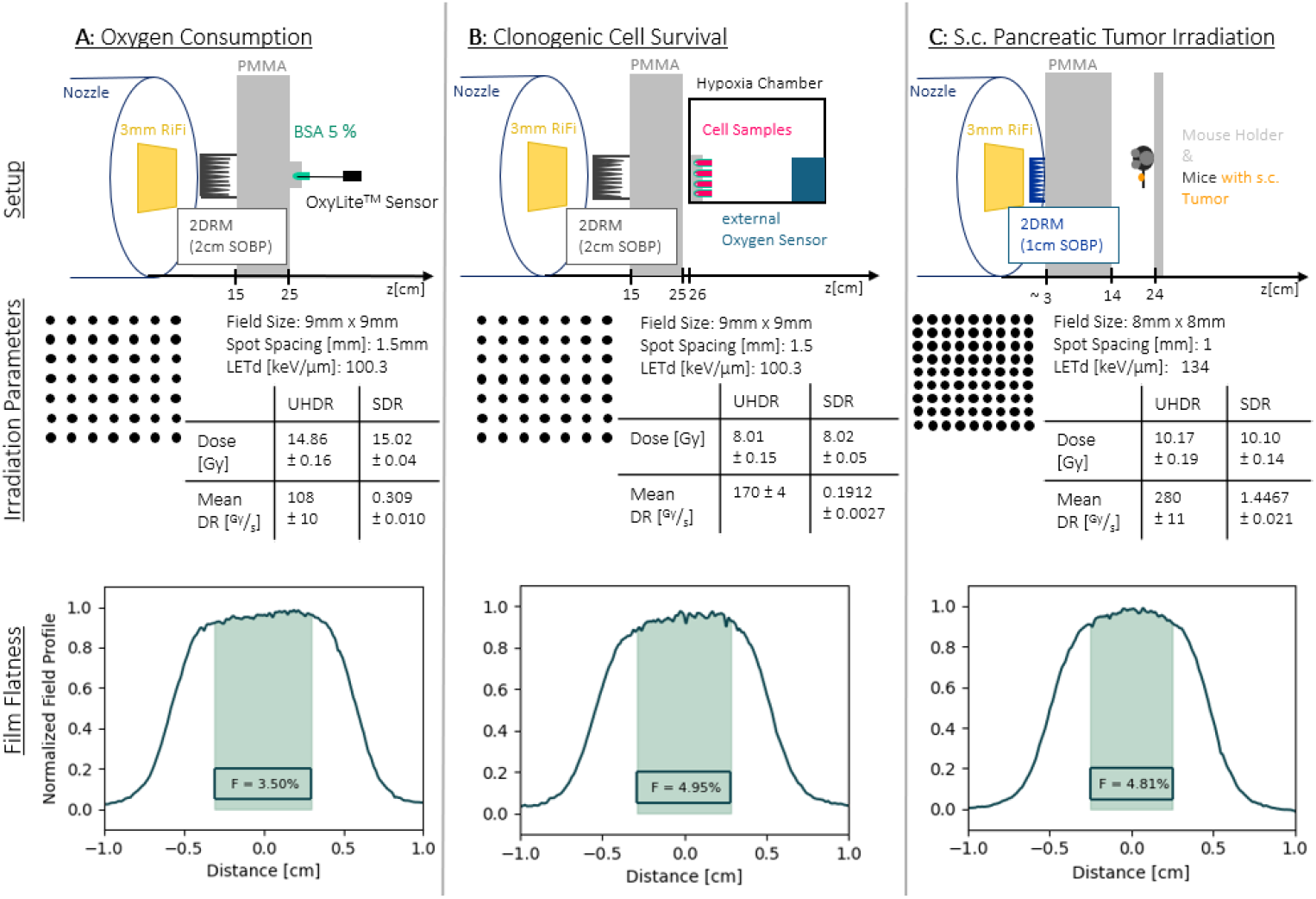
The various setups for the three experiments: A: Oxygen Consumption measurements, B: Clonogenic Cell Survival and C: S.c. Pancreatic Tumor Irradiation. The Setup illustrations contain the setup components and the spatial information. The Irradiation Parameters include details about the field sizes with the corresponding spot spacing, average LETd, dose and mean dose rate (Mean DR). The dosimetric quantities are given with their standard deviation. The Film Flatness graphs provide a representative line profile of an irradiated film with the calculated flatness index (F). As the index does not exceed 5%, homogeneous irradiation of the samples is ensured. The 3mm RiFi abbreviates the ripple filter installed in the nozzle. 2D range modulators (2DRM) producing a 1cm or 2cm spread out BP (SOBP) were used for the in vivo or oxygen consumption and in vitro experiment respectively [22], [23]. PMMA refers to Polymethylmethacrylate.

Daily dosimetric measurements were conducted with a PinPoint ionization chamber (PTW, REF: TM31015, SN:0903) following clinical practice TRS398 [21]. At least three dose measurements per experiment were conducted (Figure 1). To calculate the mean dose rate, the recordings from the Beam Application and Monitoring System (BAMS) were used to extract the length of the spills. Additionally, Gafchromic EBT3 films (8’’X10’’ (x25), Ashland 828204) were irradiated to ensure homogeneity within the fields. Information about the evaluation of irradiated films to assess the homogeneity of the fields and additional information about the spill structure can be found in the supplementary material.

### Oxygen Consumption Measurements

The effect of oxygen ion UHDR and SDR irradiation on the inherent oxygen concentration in sealed samples containing 5% BSA was examined. As reported by Karle et al [24] the oxygen consumption rate, ***g*** was calculated for each irradiation. The resulting ***g***-values were fitted using the following equation:

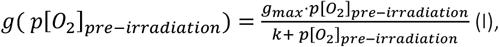

Here, p[O_2_]_pre-irradiation_ is the oxygen concentration pre-irradiation, while the fit parameters g_max_ and k correspond to the saturating g-value and to the concentration where half of the g_max_ is reached, respectively. [24] For each dose rate, three different samples were irradiated, the g-values calculated, and the data fitted with equation (I). The difference between the fit of the SDR and the UHDR data was analyzed, calculated and investigated.

### Cell culture and irradiation setup

Murine pancreatic cancer cells derived from autochthonous KPC mouse model of PDAC (KPC; 6419c5 Kerafast) [19] [20] were cultured in Dulbecco modified Eagle medium (DMEM, ATCC) supplemented with 10% heat inactivated fetal bovine serum (FBS; Gibco) and 1% penicillin/streptomycin at 37°C in a humidified atmosphere of 5% CO_2_.

For the irradiation cells were prepared as previously described [25]. The cells were placed in a hypoxia chamber (Biospherix) at ∼ 1.1% O_2_ for 5 hours pre-irradiation and maintained in hypoxic conditions during irradiation (hypoxic coindition) or at 20% O_2_ pre-irradiation and during irradiation (normoxic condition), using 8 Gy dose for both SDR and UHDR at an average LETd of 100.3keV/µm (Figure 1 B).

### Colony formation assay

Irradiated and control cells were seeded in 25-cm^2^ flasks in triplicates and incubated at 37°C at 5% CO_2_. After colony formation, cells were fixed with 75% methanol and 25% acetic acid for 10 minutes at room temperature and stained with 0.1% crystal violet for 10 minutes. Colonies containing more than 50 cells were counted as survivors. Three independent biological experiments for hypoxia and two for normoxia were conducted, each with at least three to five replicates.

The significance of difference in clonogenic cell survival between UHDR and SDR irradiation was evaluated with a Mann-Whitney-U-Test with GraphPad Prism software.

### Animal model

Subcutaneous (s.c) KPC mouse model was established by injecting 5x10^5^ KPC cells resuspended in 100µl PBS s.c. into the hind limb of 6-8 weeks-old female C57BL/6 mice (Janvier Labs). When the tumor size reached 80 ± 20 mm^3^, the mice were randomized for irradiation (n=6) and irradiated with 10 Gy SDR or UHDR oxygen ion beams at an average LETd of 134keV/µm. The mice were positioned in custom made holders and subcutaneous tumors were irradiated assuring complete tumor target coverage (Figure 1c). The tumor sizes were measured 3 times per week using a caliper. The volumes were calculated with the following formula: 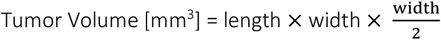.

Animal work was carried out in accordance with the rules approved by the local and governmental animal care committee established by the German government (Regierungspraesidium, Karlsruhe).

Tumor volumes were compared at given time points using one-way analysis of variance (ANOVA) with Tukey’s multiple comparison test in GraphPad Prism Software. Kaplan-Meier survival curves were compared using Log-Rank test (GraphPad Prism).

## Results

The oxygen consumption rate saturates for high initial oxygen concentrations after UHDR irradiation at 0.0205±0.0003^%^/_Gy_ and for SDR at 0.0235±0.0003^%^/_Gy_ (Figure 2, A-I). As shown in Figure 2 (A-II), the oxygen consumption rate difference between the SDR and UHDR irradiated samples saturated for high initial oxygen concentrations at 0.003^%^/_Gy_, while for the hypoxia condition the discrepancy was 0.0012^%^/_Gy_. Following the oxygen depletion hypothesis, the minimal difference in oxygen consumption observed between UHDR and SDR is unlikely to result in a measurable sparing effect in vitro. It is important to acknowledge that BSA does not fully replicate the complexity of the intracellular milieu. Therefore, this method cannot definitively elucidate an UHDR sparing mechanism but instead provides an indication of the negligible difference in oxygen consumption rates between UHDR and SDR for high-LETd irradiation at 100.3 keV/µm.

**Figure 2:**
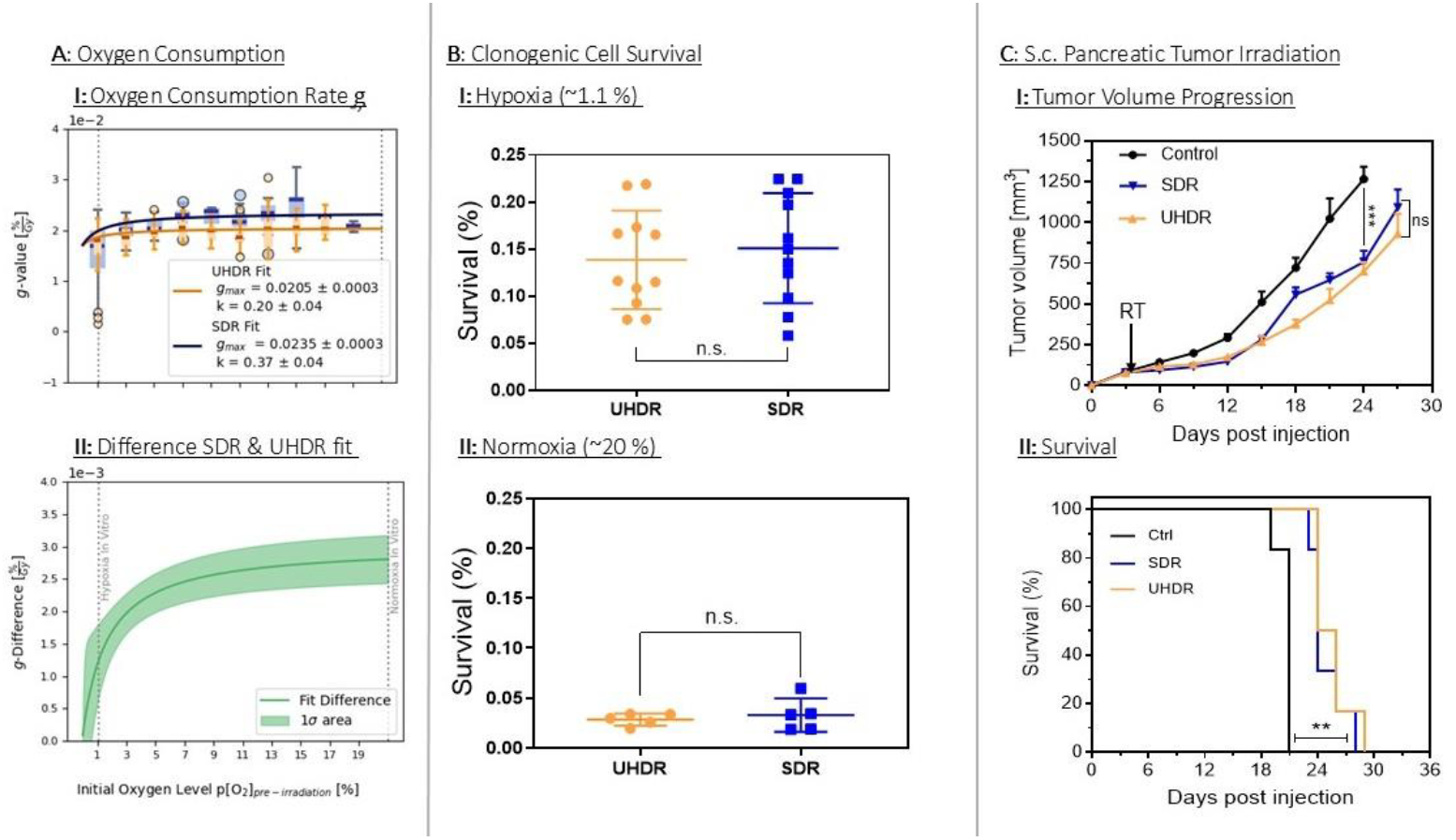
Results from the three set up are shown. (A) Oxygen consumption: (A-I) Individual g-values plotted as box plots with 2% bin intervals, displaying medians, quartiles (boxes), whiskers (extremes), and outliers. The fit functions from equation (I) are displayed as solid lines (orange for UHDR and blue for SDR) with the parameters and their standard deviation given in the legend. (A-II) Difference between UHDR and SDR g-value fits is shown as function of the initial oxygen level with standard deviation shading. In both (A-I) and (A-II) the vertical lines mark the oxygen concentration which is used in the in vitro experiment. (B) Clonogenic cell survival: Box plots show means (horizontal lines) and whiskers, with p-values calculated using the Mann-Whitney-U-Test ^n.s^p = 0.6063 and ^n.s^p = 0.8182 for hypoxia (B-I) and normoxia (B-II) respectively. (C)Tumor control and Kaplan-Meier survival: (C-I) Tumor volumes are presented as mean ± standard error of mean (N=6), with p-values from one-way ANOVA and Tukey’s post-test (^n.s.^p=0.798; ***p=0.0003). (C-II) Kaplan-Meier survival curves indicate comparable efficacy of UHDR and SDR in prolonging survival versus controls (Log-Rank test: Ctrl vs. SDR p=0.0012; Ctrl vs. UHDR p=0.012; SDR vs. UHDR p=0.3970; N=6).

Our findings with high LET oxygen ion irradiation indicate a similar cytotoxic efficacy of SDR and UHDR in normoxic and hypoxic condition. Previous in vitro studies have reported an UHDR sparing effect under hypoxia across different irradiation modalities and cell lines [25], [26], [27], [28], for doses higher than the 8 Gy, used in this work. Additionally, the LETd values in those experiments were relatively low, with carbon ions exhibiting an LETd of 13 keV/µm [27] and helium ions in the SOBP an LETd of 16 keV/µm [25]. In contrast, our study uniquely explores high LET regions (up to 100.3 keV/µm), marking the first investigation of high LET UHDR in vitro.

In vivo, both SDR and UHDR irradiations at 10 Gy single fraction and LETd 134 keV/um show significant potential for resistant KPC tumor treatment. Tumor volume reduction was substantial for both UHDR and SDR compared to controls (p=0.0003; Figure. 2, C-I). Median survival for UHDR and SDR treated animals was 24 and 25 days, respectively, compared to 20 days for the controls (p=0.0012; Figure. 2, C-II). Given the highly resistant and hypoxic nature of KPC tumor model, this extended survival is a valuable starting point for further preclinical high LET radiotherapy investigations aimed at PDAC tumor eradication.

## Conclusion

This data marks the first successful application of oxygen ion UHDR irradiation in vitro and in vivo. In terms of oxygen consumption, the difference between UHDR and SDR was found to be marginal. Elevating the biological complexity, both SDR and UHDR achieved comparable outcomes in cell survival in vitro and tumor response in vivo. The use of oxygen ion SOBP radiotherapy, characterized by its high LET (> 100 keV/µm) and superior dose conformity, demonstrates potential in addressing the challenge of treating radio resistant, hypoxic tumors, such as pancreatic cancer. These results underscore the potential of oxygen ion therapy to address the challenges of pancreatic tumor treatment by offering more precise dose distribution with high LET. Additionally, oxygen ion beams at UHDR could potentially minimize motion-related uncertainties.

## Acknowledgement

The authors would like to thank the whole accelerator team at HIT for helping with the synchrotron adjustments and Uli Weber from Technische Hochschule Mittelhessen Campus Giessen for providing the 2DRMs.

