## Supplemental Material for "Exploring High LET Oxygen-Ion FLASH Radiation for Targeting Pancreatic Cancer in Vitro and In Vivo"

### Supplementary Material

#### Irradiation facility

The Heidelberg Ion Beam Therapy Centre (HIT) currently facilitates research using four particle beams: protons, helium, carbon and oxygen ions. The latter two are generated using a single electron cyclotron resonance ion source, with carbon dioxide ($CO_2$) as primary gas, yielding carbon ions ($^{12}C_4$+) or oxygen ions ($^{16}O_6$+) [1]. After acceleration in a synchrotron, the particles are extracted via a third-order resonant RF-knockout extraction system. The nozzle is equipped with a Beam Application and Monitoring System (BAMS), which integrates various ionization chambers and multi-wire proportional chambers. These components serve the dual purpose of monitoring and recording the spill structure to facilitate the raster scanning. Additionally, this feedback loop enables precise intensity-controlled application of raster-scanned pencil beams [2].

To achieve UHDR, specific synchrotron adaptations are required. These include tuning the extraction frequency closer to resonance and increasing sextupole magnet strength. Notably, these modifications facilitate UHDR application without compromising the functionality of the raster scanning or intensity control systems [3].

#### Spill Structure Information

| Experiment | Field Size [$mm^2$] | Spot Spacing [mm] | Mean LETd [keV/µm] | Mean Dose Rate [Gy/s] | Delivery Time [s] | Spill Time [ms] | Interspill Time [s] | Dose [Gy] |
|---|---|---|---|---|---|---|---|---|
| Oxygen Consumption | 9 x 9 | 1.5 | 100.3 [range 88.0–235.0] | 0.309 ± 0.010 | 48.5 ± 1.6 | 876 ± 154 | 4.39 ± 0.05 | 15.02 ± 0.04 |
| | | | | 108 ± 10 | 0.138 ± 0.012 | 137 ± 12 | - | 14.86 ± 0.16 |
| In Vitro | 9 x 9 | 1.5 | 100.3 [range 88.0–235.0] | 0.1912 ± 0.0027 | 41.8 ± 1.3 | 367 ± 9 | 4.23 ± 0.04 | 8.02 ± 0.05 |
| | | | | 170 ± 4 | 0.047 ± 0.003 | 47 ± 3 | - | 8.01 ± 0.15 |
| In Vivo | 8 x 8 | 1 | 134 [range 106–162] | 1.4467 ± 0.021 | 6.98 ± 0.03 | 1631.8 ± 99 | 4.3 ± 0.04 | 10.10 ± 0.14 |
| | | | | 280 ±11 | 0.0361 ± 0.0013 | 36.1 ± 1.3 | - | 10.17 ± 0.19 |

*Table 1: Spill structure information for all three experiments. The mean dose rate is calculated by dividing the total dose applied by the total irradiation time. The delivery time encompasses the interval between the initiation of the first pulse and the conclusion of the final pulse, whereas the spill time represents the temporal duration of a single spill. The interspill time, on the other hand, denotes the interval between the end of one pulse and the subsequent onset of the next.*

#### Film Evaluation

To guarantee that the various field sizes and spot spacings employed in the active pencil beam scanning irradiation deliver a uniform dose to the samples, EBT3 Gafchromic Films (Ashland) were irradiated and scanned at a resolution of 1200 dpi. Horizontal and vertical line profiles through the center of the film were extracted with ImageJ. In python, the profile was normalized to the maximum greyscale value and smoothed with a Gaussian filter. From the central point of the profile, we identified a region spanning 50% of the total area under the curve. Within this central region, we calculated a flatness index (F):

$$F = 100 * \frac{Y_{max} - Y_{min}}{Y_{max} + Y_{min}}$$

with Ymax and Ymin being the maximum and minimum normalized grey scale value in the central area. The profile was evaluated for one film per experiment and the profile with the largest F was displayed in Figure 1.
